# Exploiting the Angiotensin-Converting Enzyme Pathway to Augment Endogenous Opioid Signaling

**DOI:** 10.1101/2025.02.19.639161

**Authors:** Prakashkumar Dobariya, Jessica Williams, Filip Hanak, Patrick E. Rothwell, Swati S. More

**Author notes:** Corresponding Author: Swati S. More.

## Abstract

Angiotensin Converting Enzyme (ACE) impacts hemodynamics by regulating the conversion of angiotensin I to the vasoconstricting angiotensin II. We recently identified *a non-canonical central* role of ACE in the degradation of enkephalin heptapeptide, Met-enkephalin-Arg-Phe (MERF). Enkephalins are short-lived, endogenous opioid peptides that mediate the body’s intrinsic analgesic response. Here we identify chemically diverse ACE inhibitors using an optimized high throughput screening assay to boost endogenous opioid signaling. Our primary hits (thiorphan, D609, and raloxifene) were selected for dose-response characterization, *in vitro* enkephalin release, *in vivo* analgesic potency, and *in silico* analysis. Intracerebroventricular administration of these compounds significantly attenuated pain response, alone and in combination with MERF, which was reversed by opioid receptor antagonist naloxone. Molecular docking provided additional insight into the active site interactions of these scaffolds, which could be exploited further for creation of more potent inhibitors. These results showcase the potential of central ACE inhibitors to modulate endogenous MERF signalling.

**Graphical Abstract:** 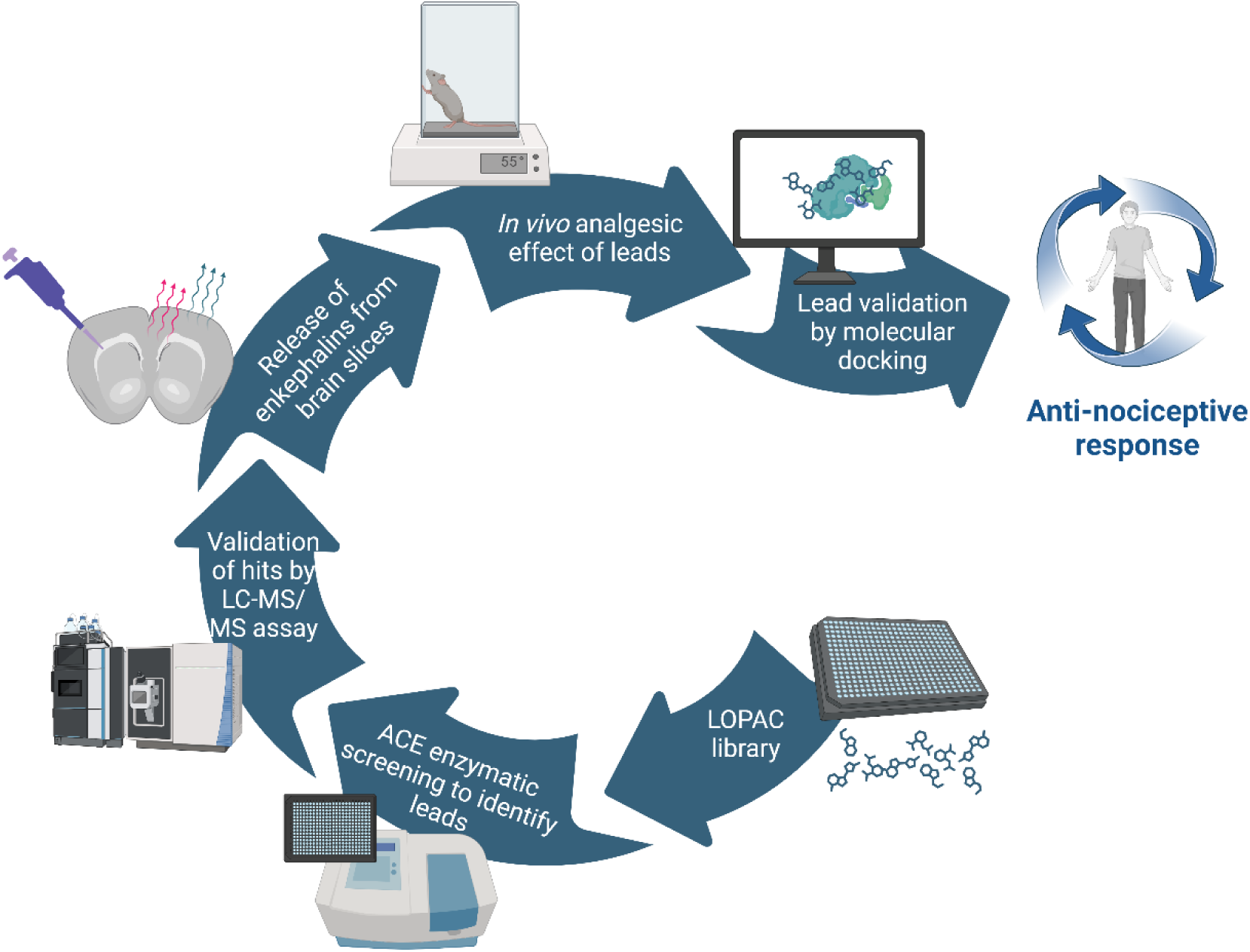

## Introduction

The Renin-Angiotensin system (RAS) is a multifaceted system that regulates diverse physiological processes. The system is characterized by two opposing pathways: the Angiotensin Converting Enzyme (ACE)-Angiotensin II (Ang II)-AT1 receptor axis, which exerts vasoconstrictive and pro-inflammatory effects, and the ACE2-Angiotensin (1-7)-MAS receptor axis, which counteracts these effects through vasodilatory and anti-inflammatory actions. The components of these pathways are expressed throughout all tissue and organs, forming a localized angiotensin system characteristic of each area [1]. ACE enzyme, a key component of the RAS, is responsible for converting angiotensin I (Ang I) to Ang II, involved in the regulation of the cardiovascular system, and has found application in the development of clinically used antihypertensive medications. Beyond the role in the regulation of blood pressure, electrolyte/fluid balance, and other circulating humoral system, we recently reported a non-canonical role of ACE in the regulation of endogenous opioid peptides, termed enkephalins (ENKs) [2, 3]. The study demonstrated that while ACE can cleave enkephalin peptides, it is not principally responsible for the degradation of conventional Met-enkephalin or Leu-enkephalin in brain tissue. It is involved in the degradation of the heptapeptide, Met-Enkephalin-Arg-Phe (MERF), which is highly expressed in the nucleus accumbens and displays a higher binding affinity for the μ-opioid receptor. The nucleus accumbens is a central hub in brain reward circuits impacted by drugs of abuse. Thus, inhibition of ACE by captopril enhanced extracellular levels of MERF, resulting in cell type-specific depression of excitatory synaptic transmission due to the activation of opioid receptors, and attenuating conditioned place preference produced by fentanyl [3].

Enkephalins such as MERF were discovered in 1975 by John Hughes, and are found to be widely distributed throughout the body, including the central and peripheral nervous systems, immune cells, and adrenal glands. In the brain, enkephalins are abundant in the rostral ventromedial medulla, hypothalamus, amygdala, and periaqueductal grey [4, 5]. These peptides are synthesized from the precursor peptide proenkephalin (PENK), whose cleavage by enkephalin-degrading enzymes (enkephalinases) liberates peptides of varying length and sequence such as the pentapeptides, Met-enkephalin (Tyr-Gly-Gly-Phe-Met) and Leu-enkephalin (Tyr-Gly-Gly-Phe-Leu), heptapeptide MERF (Met-enkephalin-Arg-Phe), and octapeptide MERGL (Met-enkephalin-Arg-Gly-Leu). Tyr and Phe are required for their binding to the opioid receptors. Enkephalins mediate their effect through activation of the µ-opioid receptor (MOR) and δ-opioid receptor (DOR) of spinal and supraspinal sites with differential selectivity [6], and are involved in the regulation of motor control, feeding behavior, emotions, and pain [7–9]. Enkephalins interrupt nociceptive signal transmission at the dorsal horn by suppressing the release of neuropeptides such as substance P, glutamate, and calcitonin gene-related peptide (CGRP), which are involved in the transmission of pain signals. Additionally, enkephalins also control glutaminergic, serotonergic, and noradrenergic neurons and resultant release of glutamine, serotonin and norepinephrine that inhibit pain transmission [10, 11]. Thus, enkephalins regulate the excitatory and inhibitory signals within the pain pathways, ultimately leading to a reduction in pain perception [12]. Previously, enkephalins and their analogs have been studied for their analgesic response in animal models of pain, specifically for their reduced potential for respiratory depression and dependence, commonly observed with exogenous opioids [13]. The pharmacological action of enkephalins is compromised due to their rapid degradation by peptidases. Thus, increasing endogenous levels of these opioid peptides by inhibiting enkephalinases could present an alternative effective strategy for mitigating pain [13, 14]. Given the short half-life of enkephalins, preventing their degradation by such peptidase inhibitors could avoid overstimulation of the opioid receptors and thus prevent the development of dependence.

With the recently identified role of ACE in the degradation of potent enkephalin peptide MERF, specific inhibitors of ACE could increase MERF levels and enhance the body’s natural pain-relief mechanisms. In fact, combining ACE inhibitors with opioids can present synergistic analgesia. Captopril and other ACE inhibitors potentiate morphine-induced analgesia [15–17]. When given in a sub-analgesic dose, captopril in combination with the sub-analgesic dose of MERF has been shown to double jumping latency in response to thermal stimulus compared to when given alone [18]. These studies suggest that targeting the ACE-enkephalin axis may offer a promising therapeutic approach to pain management. ACE is a pleotropic enzyme, its differential physiological functions and effects are dependent on the distinct catalytic domains. The somatic isoform of ACE (sACE) consists of two catalytic domains (N- and C-domains) with active site zinc-binding motif HEMGH (His-Glu-Met-Gly-His); while ACE expressed in sperm cells, termed testicular ACE (tACE), comprises of only one catalytic domain identical to the C-domain. The 45% sequence divergence between the N- and C-domains of ACE results in their distinct kinetic properties and substrate specificity, suggesting functional differences between the two domains [19]. For instance, the N-domain of the ACE is more effective in the hydrolysis of angiotensin 1-7, enkephalins, kinins, neurotensin, formyl-Met-Leu-Phe, substance P, gonadotropin-releasing hormone (GnRH), and N-acetyl-Ser-Asp-Lys-Pro (AcSDKP) [20, 21]. Distinct substrate specificities of the N- and C-domains of ACE offer a promising approach for designing domain-specific, structure-based drug entities, enabling the development of targeted therapies that can selectively modulate specific ACE functions. The utility of domain selective inhibitors has been demonstrated in the design of anti-hypertensives, in which complete inhibition by a non-selective inhibitor resulted in side effects due to the excessive levels of bradykinin. In contrast, the use of C-domain preferring or selective inhibitors helped to reduce blood pressure without affecting the N-domain selective functions [22, 23]. Although the applications of the domain-selective inhibitors have been reported with other physiological functions, their role in pain is poorly explored. An *in vitro* study using purified ACE has indicated the involvement of ACE N-domain in the degradation of MERF [24, 25]. Thus, selective targeting of the N-domain may offer a promising avenue for developing effective analgesics without or minimally affecting the cardiovascular function of ACE.

In this study, we seek to identify chemically diverse ACE inhibitors by employing high throughput screening (HTS) of a commercially available library of bioactive small molecules. We utilized a miniaturized fluorescence-based assay in a 384-well plate and screened a Library of Pharmacologically Active Compounds (LOPAC) comprising of over 1,200 compounds in an enzymatic assay utilizing somatic ACE. Several promising candidates were subsequently analyzed for their half-maximum inhibitory concentrations (IC_50_) using ACE domain-specific and non-domain-specific substrates to evaluate their domain selectivity. The antinociceptive effect of the lead compounds were evaluated in tail flick and hot plate assays, alone and in combination with MERF. Finally, computational studies were conducted to elucidate their interactions with the target and to gain insight into future structure exploration studies for improving ACE inhibitory potency. Collectively, this study reports optimization of FRET-based ACE enzymatic assay for HTS application to identify chemically diverse inhibitors for modulation of endogenous opioid signaling. The results provide the proof-of-concept in support of the development of domain-selective ACE inhibitors as single agents or adjuvants to opioids for clinical management of pain.

## Results

### Optimization of the HTS enzymatic assay

Numerous assays for measuring pharmacological inhibition of ACE have been reported previously. These methods include spectrophotometric [26, 27], fluorometric [28], radiochemical [29], high-performance liquid chromatography with a UV detector (HPLC-UV) [30], capillary electrophoresis [31], and mass spectrometric methods [32]. These assays involve use of different peptide substrates such as hippuryl-l-histidyl-l-leucine (HHL) [27, 33], o-aminobenzoylglycyl-p-nitrophenylalanyl proline [28], furanacryloyl-l-phenylalanylglycyl-glycine (FA-PGG) [34] and 3-hydroxybutyryl-glycyl-glycylglycine (3-HB-GGG) [35]. While these methods are effective for evaluating ACE activity, they may not be ideal for high-throughput screening due to factors such as sample preparation time, reagent costs, and analytical complexity. We aimed to develop a rapid, accurate, and reproducible high-throughput fluorescence-based method for screening potential ACE inhibitors. We selected Mca-RPPGFSAFK(Dnp)-OH as a fluorescence substrate over previously reported weak fluorescent substrate (o-aminobenzoylglycyl-p-nitro-L-phenylalanyl-L-proline) for this enzymatic reaction. The substrate consists of a highly fluorescent 7-methoxycoumarin group, which is internally quenched by 2,4-dinitrophenyl group (Figure 1A). The release of 7-methoxycoumarin-4-acetyl (Mca) upon cleavage by ACE and resultant increment in fluorescence was monitored by excitation at 320 nm and emission at 405 nm for 30 min. A typical reaction kinetics is shown in the Figure 1B. The affinity of ACE for the substrate was calculated from the Michaelis-Menten equation under these experimental conditions, and provided a *K*_m_ of 14 ± 0.74 μM (Figure 1C).

**Figure 1.**
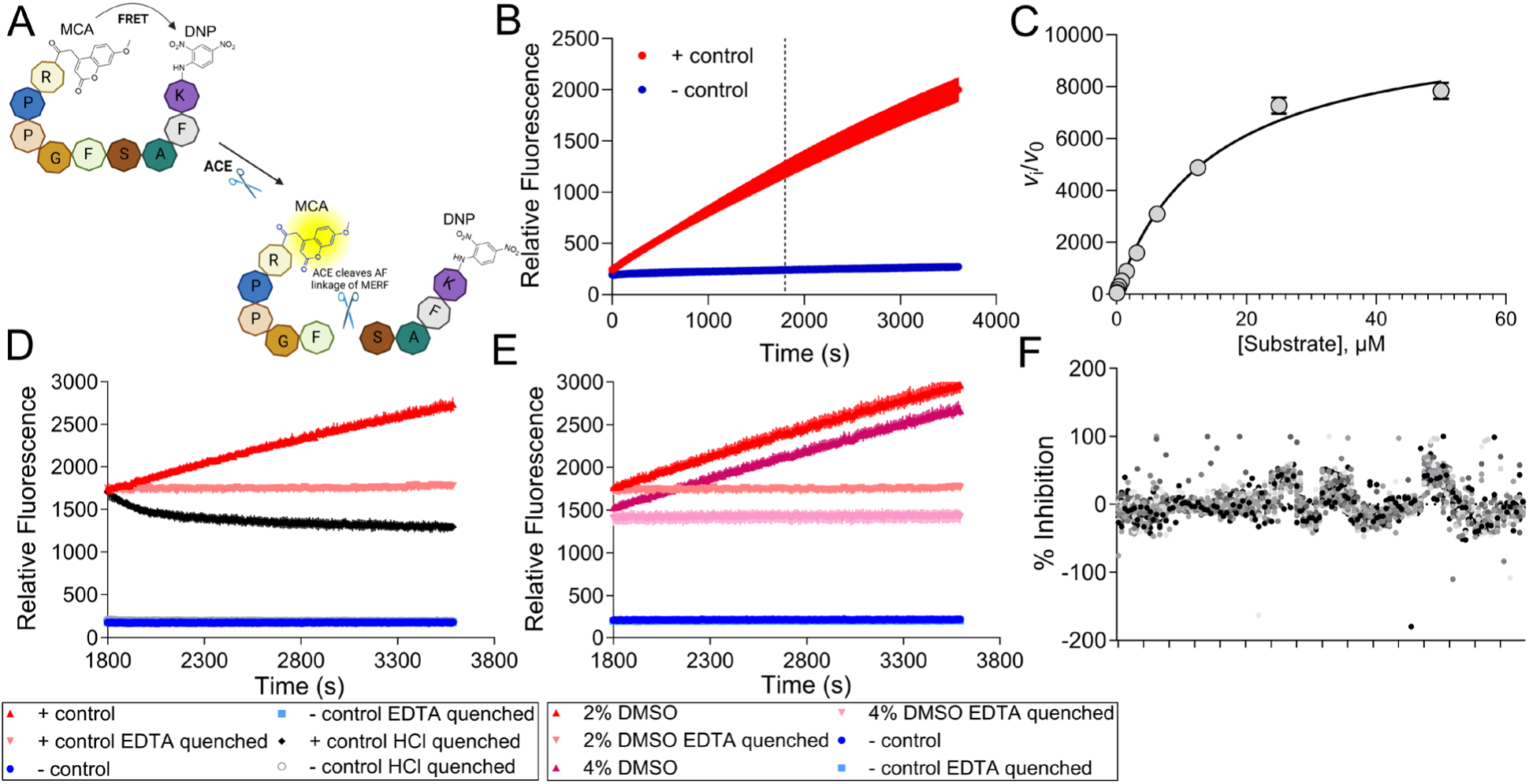
Optimization of ACE enzymatic assay for high throughput screening of the LOPAC library using a fluorescent peptide substrate. (A) Schematic of the fluorescence assay using internally quenched Mca-RPPGFSAFK(Dnp)-OH peptide, which upon cleavage by mACE releases fluorescent MCA. Amino acid residues are indicated with a single letter (Abbreviations: Mca – 7-Methoxycoumarin-4-yl-acetyl; Dnp - 2,4 Dinitrophenyl, ACE-angiotensin converting enzyme). (B) A time course of a typical ACE reaction using the fluorescent peptide under conditions stated in Methods. A linear reaction progress was noted until 1 h. The vertical line shows the time point at which the reaction was quenched in D and E. (C) The Michaelis-Menten kinetics of the ACE reaction was performed with increasing concentrations of the fluorescent substrate. The initial velocity (*V*_i_) of the reaction was calculated as described in Methods and a graph was plotted to calculate *K*_m_ for the substrate. (D) Exploration of EDTA and HCl as reaction quenchers, added at 30 minutes after the substrate addition, and continued for additional 30 minutes. (E) Assessing DMSO tolerance of the fluorescence assay by conducting the reaction in the presence of 2% and 4% DMSO, followed by a quenching step with EDTA. (F) Scatter plot illustrating the percent inhibitory activities of the test compounds from the LOPAC library.

To facilitate the assay application for high throughput screening, we optimized several of the assay parameters. A quenching step was incorporated at the end of the enzymatic reaction, which included either decreasing the reaction pH (HCl) or metal chelation (EDTA). After the addition of 1 N HCl or 4 mM EDTA at the end of the 30-minute incubation period, the reaction was continued for an additional 30 minutes to determine the effect of these quenchers on the fluorescence signal. As shown in Figure 1D, quenching with 1 N HCl resulted in a slight decrease in the fluorescence intensity, while EDTA quenching led to a stable fluorescence signal, which was unchanged during the additional incubation period.

Given that the library compounds are dissolved in DMSO, we examined the effect of DMSO concentration on enzyme activity. We found that ACE activity was unaffected even at 4% DMSO (Figure 1E). Further, we included a reducing agent TCEP (1 mM) and a surfactant IGEPAL (0.0025%) in the assay buffer to maintain the correct protein structure/activity, and to prevent false positives resulting from compound aggregation, respectively. Based on these findings, the final assay conditions for our screening consist of 7.5 ng of the enzyme, 10 μM of the substrate peptide, and assay buffer containing a final concentration of DMSO at 3.3%, TCEP at 1 mM and IGEPAL at 0.0025% in 50 mM MES (pH 6.5).

### HTS of a commercial library of small molecules

This optimized HTS protocol was employed to screen a commercially available library of pharmacologically active compounds (LOPAC) containing 1,280 small molecules. The robustness of the optimized assay was confirmed by a Z’ factor of 0.76, calculated from 48 positive and 48 negative controls. A fixed concentration (41.7 μM) of the compounds was utilized for initial screening. The mean percentage inhibition of the library compounds was 4% with a standard deviation of 13.9% and is displayed as a scatter plot in Figure 1F. For identification of potential hits from the screen, we set a hit threshold of ≥ 46% inhibition based on the mean percentage inhibition plus three standard deviations as described in Methods. A total of 19 compounds were identified as hits, offering a hit rate of 1.5% (Supporting Information, Table S1). The screen also identified three known ACE inhibitors, captopril, trandolapril, and enalaprilat dihydrate, as hits in our assay, providing further validation of our experimental approach.

### Dose-response studies of the lead compounds against mACE

To further confirm and investigate dose dependency of the inhibitory effects, the compounds were rank ordered based on inhibitory potency (Supporting Information, Table S1), and six of the leads (SKF 89976A, thiorphan, lonidamine, D609, raloxifene) along with captopril were selected for dose-response studies based on their commercial availability and potential central effects. Compounds SKF 89976A and lonidamine did not confirm the HTS findings and were not considered for further validation studies. Various concentrations of the remaining three inhibitors (Figure 2A) were incubated in the enzymatic reaction, which displayed competitive sigmoidal dose-dependent inhibition. The enzymatic curves of the velocity versus inhibitor concentrations were utilized to calculate IC_50_ values (Figure 2B). Captopril, a known potent ACE inhibitor, offered the lowest IC_50_ of 0.013 ± 0.002 µM, which is in agreement with previously reported values of 1-20 nM [36, 37]. Thiorphan, D609, and raloxifene offered IC_50_ values of 2.36 ± 0.42 µM, 31.05 ± 10.82 µM, and 102.24 ± 13.23 µM, respectively (Table 1).

**Figure 2.**
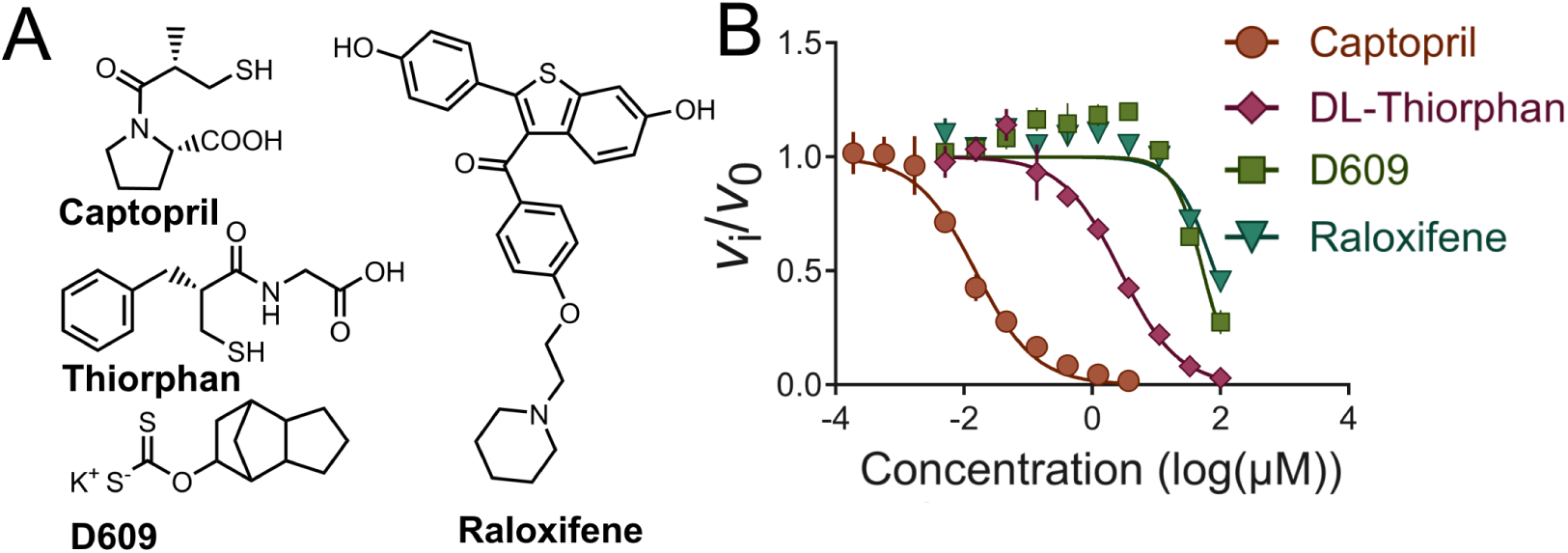
Dose-response analysis and determination of IC_50_ values of the selected lead compounds. (A) Chemical structures of the identified LOPAC hits — captopril (positive control), thiorphan, D609, and raloxifene hydrochloride. (B) Dose-response curves of the compounds obtained by fluorescence-based ACE enzymatic assay. The calculated IC_50_ values of the lead compounds from the fluorescence and mass spectrometry-based methods are provided in Table 1.

**TABLE 1:** Calculated IC_50_s of the identified hits from the high-throughput screen.

| Compound | $IC_{50} \pm SEM$ (μM)<br>by fluorescence | $IC_{50} \pm SEM$ (μM)<br>by LC-MS/MS |
| --- | --- | --- |
| Captopril | $0.0130 \pm 0.002$ | $0.006 \pm 0.001$ |
| Thiorphan | $2.360 \pm 0.42$ | $3.510 \pm 0.37$ |
| D609 | $31.05 \pm 10.82$ | $44.90 \pm 5.21$ |
| Raloxifene | $102.24 \pm 13.23$ | $88.57 \pm 3.12$ |

### Confirmation of Selected Hits by LC-MS/MS assay

To confirm the inhibitory effects of the lead compounds, we utilized a secondary assay based on mass spectrometry. Herein, we measured the relative amounts of intact residual substrate peptide, Mca-RPPGFSAFK(Dnp)-OH remaining at the end of the enzymatic reaction by LC-MS/MS. The instability of the substrate peptide in the assay mixture after acetonitrile-mediated protein precipitation and its low solubility posed challenges. Preparation and analysis of the samples at room temperature and limiting the number of samples in each experimental run to 16 effectively solved such problems. The inhibitory effect of the hits was calculated using captopril as the positive control and DMSO-only reaction as the negative control. Captopril showed the lowest IC_50_ (0.006 ± 0.001 µM), which corresponded with its maximal ACE inhibitory effect. IC_50_s of the other leads, thiorphan, D609, and raloxifene, by mass spectrometry were 3.51 ± 0.37 µM, 44.90 ± 5.21 µM and 88.87 ± 3.12 µM, respectively. Individual dose-response curves for these inhibitors obtained from fluorescence and mass spectrometry-based assays are shown in Figure S1 (Supporting Information). The IC_50_ values obtained from the LC-MS/MS analysis are in close agreement with the IC_50_s obtained by the fluorescence-based ACE enzymatic assay. Collectively, our pilot screening and the subsequent secondary validation by dose-response studies and LC-MS/MS analysis demonstrate the reliability of our HTS assay for identifying potential ACE inhibitors from compound libraries.

### Measurement of enkephalin release from brain slices in the presence of lead compounds

Endogenous enkephalins exert their effects by activation of opioid receptors. However, their activity is limited by degradation by enkephalinases such as ACE. In our previous study, we showed that captopril can reduce the breakdown of endogenous enkephalins, specifically that of MERF, in experiments involving stimulation of peptide release from brain slices. We performed similar experiments in the presence and absence of the identified LOPAC leads (Figure 3A). Different concentrations of the inhibitors were employed based on their IC50s (captopril and thiorphan at 10 μM, while raloxifene and D609 at 500 μM). The release of opioid peptides was stimulated by KCl (50 mM) and the released enkephalins were measured in the extracellular fluid using our previously established LC-MS/MS method. Similar to captopril, thiorphan and raloxifene could significantly increase extracellular levels of MERF to approximately two-fold higher than the vehicle control group (Figure 3B). D609, however, was unable to demonstrate this effect on MERF release under these experimental conditions. Interestingly, the PAN enkephalinase inhibitor, thiorphan, affected the release of Met- and Leu-enkephalins, which was not observed in raloxifene-treated brain slices (Figure 3C, 3D). Given our previous observation regarding the specific role of ACE in modulating MERF levels, these results could indicate the specificity of raloxifene in offering an ACE-mediated effect on endogenous opioid signaling.

**Figure 3.**
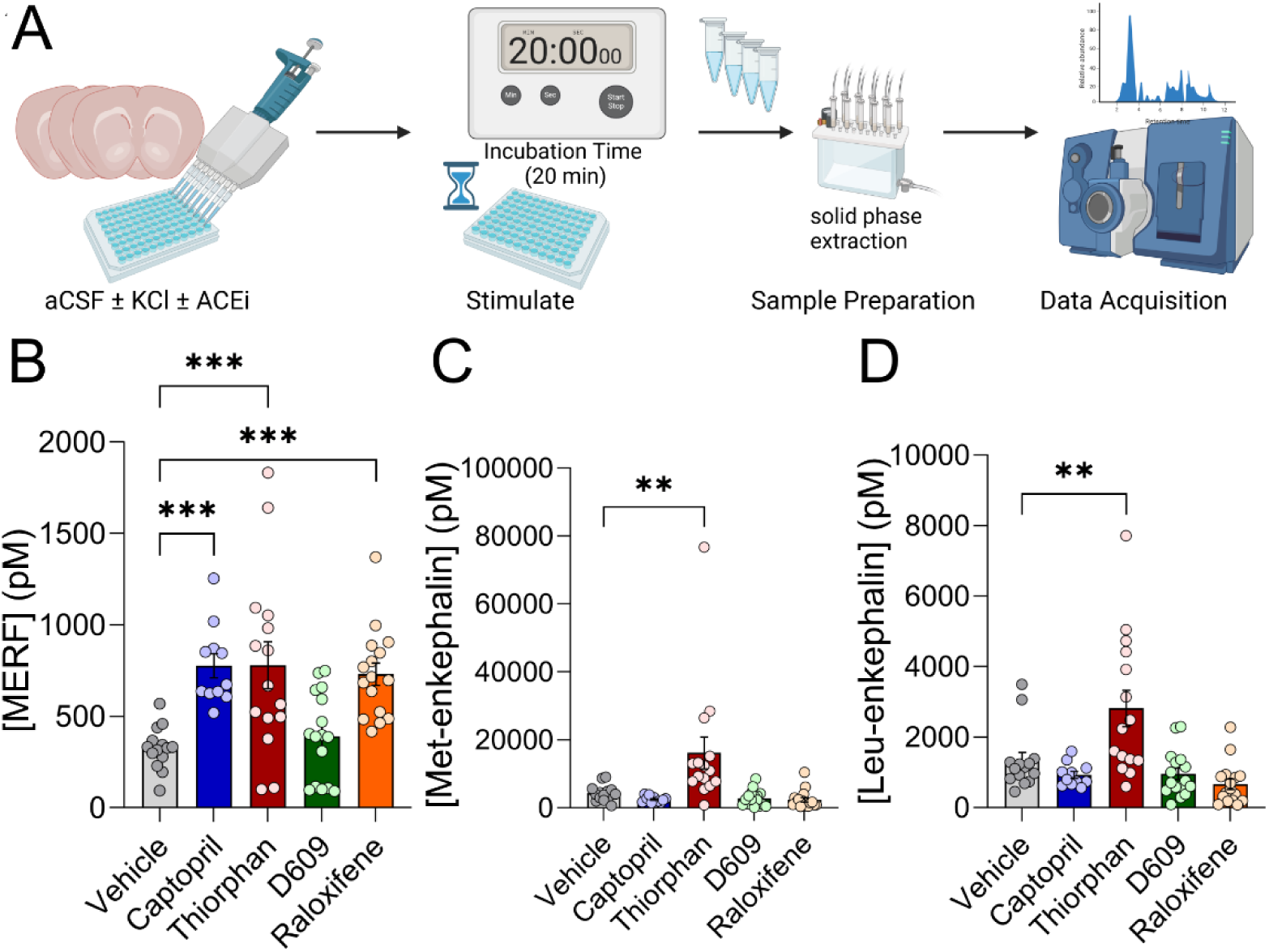
Determining the effect of ACE inhibition by the selected LOPAC leads on KCl-stimulated endogenous enkephalin release. (A) Schematic of the experimental plan. (B) LC-MS/MS analysis of enkephalins reveal a significant elevation of MERF levels by captopril, thiorphan, and raloxifene. Thiorphan also increased Met-enkephalin (C) and Leu-enkephalin (D) levels. Data are shown as the mean ± SEM. One-way ANOVA with Dunnet’s post-hoc test was used to determine the statistical significance (** *p* ≤ 0.01, *** *p* ≤ 0.001, ns = not significant) (N=6-8/treatment).

### In vivo behavior assay to assess pain perception

#### Tail flick assay

Tail-flick assay is widely used for evaluating the analgesic potency of small molecules. It quantifies the nociceptive reflex elicited by a controlled thermal stimulus applied to the tail. The leads from the HTS were subjected to analgesic efficacy evaluation *in vivo*. The compounds were administered by *i.c.v.* route to record the latency to thermal stimulus and the results are expressed as %MPE as described in Methods. The choice of the route was based on the unknown brain permeability of these compounds. The time course of the data showed the highest %MPE at 15 minutes post compound administration, with sustained effects lasting for about 45 minutes (Figure 4, upper panel). Data from the 15-minute time point was used to compare various treatment groups (Figure 4, lower panel). Compound-only groups showed significantly higher %MPE compared to the corresponding vehicle group (captopril: 21.68%, thiorphan: 22.45%, raloxifene: 13.33%, over vehicle control), although the effect is modest. In the case of D609, however, the increase in %MPE was accompanied by visibly scruffy fur and lethargy in animals. When administered at a lower 75 µg/mouse dose, D609 caused a similar increase in %MPE, confirming its modest analgesic potential (Supporting Information, Figure S2). The results of this study indicate modest potentiation of endogenous opioid signaling by these compounds leading to analgesic response.

**Figure 4.**
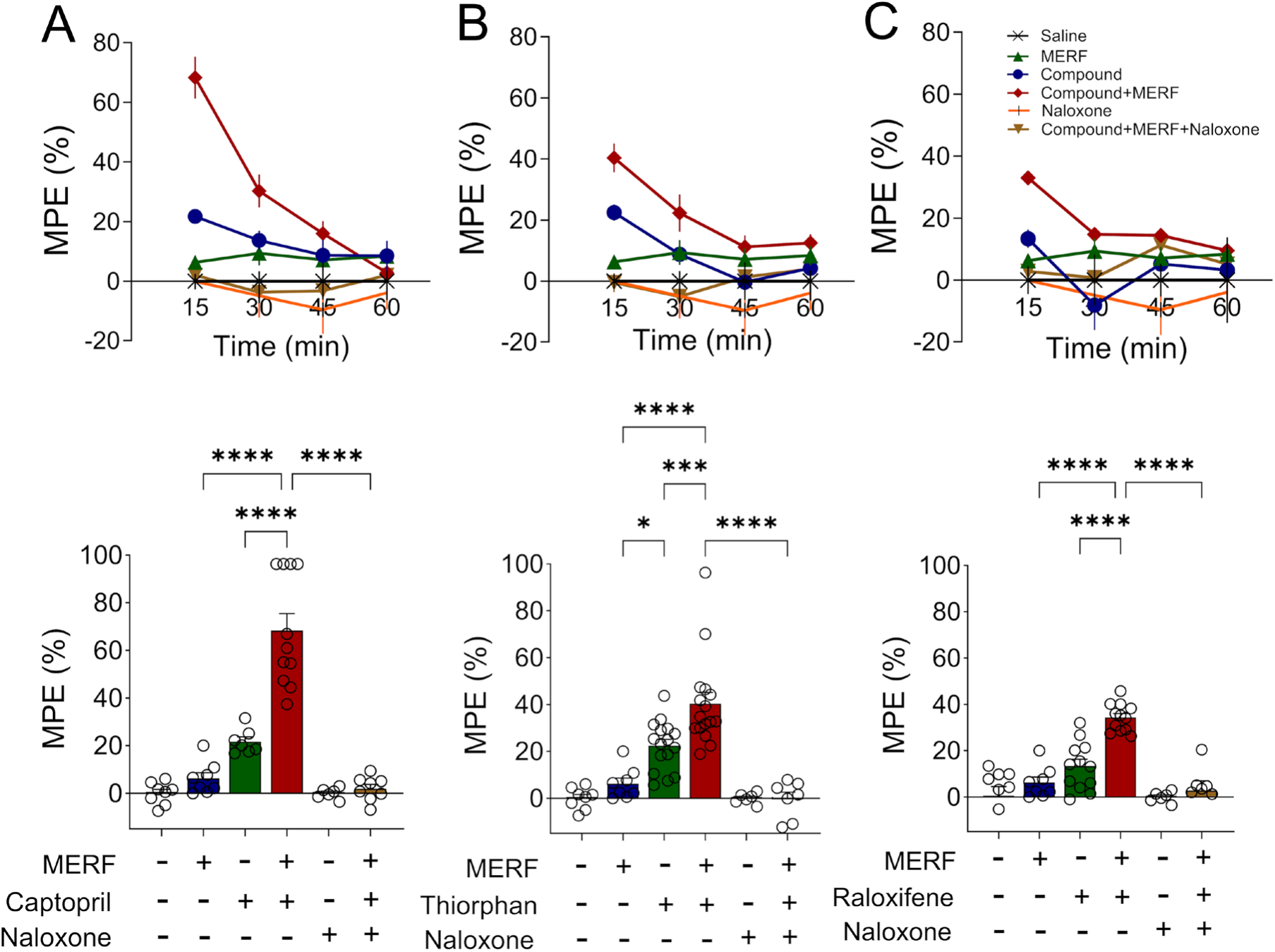
Assessment of the antinociceptive effect of the lead compounds by tail-flick assay. Compounds (100 μg/mouse) were administered directly to the mouse brain by intracerebroventricular (*i.c.v.*) route, either alone or in combination with the MERF (50 μg/mouse) and the maximum possible effect of the compounds was recorded (N = 6–15/group). Naloxone (5 mg/kg, s.c), an opioid receptor antagonist was injected to confirm the role of opioid receptor activation in the antinociceptive effect of the compounds. The top panel shows a time-course of the tail-flick response after *i.c.v* administration of captopril (A), thiorphan (B), and raloxifene (C). The Y-axis denotes the %MPE and the X-axis shows the time points at which the response was recorded. The bottom panel shows the comparison between the anti-nociceptive effect (%MPE ± SEM) for various treatment groups shown in the top panel at 15 minutes after compound administration. Data are shown as the mean ± SEM. Statistical significance was examined by a one-way ANOVA with Sidak’s post-hoc multiple comparison test (* *p* < 0.05, *** *p* < 0.001, **** *p* < 0.0001).

To further confirm the effect on enkephalin levels, the compounds were administered in combination with MERF. Exogenous supplementation of MERF did not show an analgesic response (% MPE: 6.27%), possibly due to its rapid degradation *in vivo*. The combination of captopril, thiorphan, and raloxifene with MERF showed a statistically significant increase in %MPE over each compound given alone (captopril vs captopril + MERF, 21.68% vs 68.31%, respectively; thiorphan vs thiorphan + MERF, 22.45% vs 40.32%, respectively; raloxifene vs raloxifene + MERF, 13.33% vs 34.28%, respectively). Due to pH differences and precipitation issues, the combination of D609 with MERF was administered in two separate *i.c.v.* injections. Only a marginal increase in %MPE was offered by D609 + MERF combination, compared to the changes caused by thiorphan and raloxifene combinations with MERF, suggestive of lower *in vivo* efficacy of D609 at the dose chosen to compensate for its apparent toxicity (Supporting Information, Figure S2).

The involvement of opioid receptor activation in the analgesic response was further confirmed by the administration of a non-specific opioid receptor antagonist, naloxone. Although naloxone itself is unable to induce analgesia, subcutaneous administration of naloxone after *i.c.v.* administration of the combinations (compound + MERF) resulted in attenuation of the analgesic response (Figure 4). This reversal of tail flick response by naloxone further suggests the activation of endogenous opioid signaling by ACE inhibitors.

#### Hot plate assay

To further confirm these findings, a hot plate assay was performed following a similar experimental design. Hot plate assay measures the time required by the animal to lift or shake the hind paw in response to a thermal stimulus as %MPE. The time course of the hot-plate analgesic response is shown in the upper panel of Figure 5A-C. Similar to the tail-flick assay, the peak analgesic effect was recorded at 15 minutes post-injection, with captopril exhibiting the highest %MPE. The trend of the data obtained from the hot plate assay was consistent with that of the tail-flick assay, further corroborating the compound’s ability to induce analgesia. A bar graph analysis of the 15-minute time point further emphasizes the superior analgesic efficacy of the compound-only treated groups compared to vehicle-treated controls (captopril: 10.32%, thiorphan: 13.83 %, raloxifene: 5.52 %, over vehicle control). The animals administered with combinations of these compounds with MERF showed significantly improved hot plate response over the compound-only groups (captopril vs captopril + MERF, 10.32% vs 51.30%, respectively; thiorphan vs thiorphan + MERF, 13.83% vs 36.48%, respectively; raloxifene vs raloxifene + MERF, 5.52% vs 25.89%, respectively), recapitulating the analgesia potentiation observed in the tail-flick assay. This potential synergistic interaction in analgesia between compounds and MERF was inhibited by naloxone (Figure 5), supporting the involvement of opioid receptors. The data from these acute pain models support our rationale that the leads could provide endogenous pain relief by protecting and potentiating the effects of MERF, partly by inhibiting ACE.

**Figure 5.**
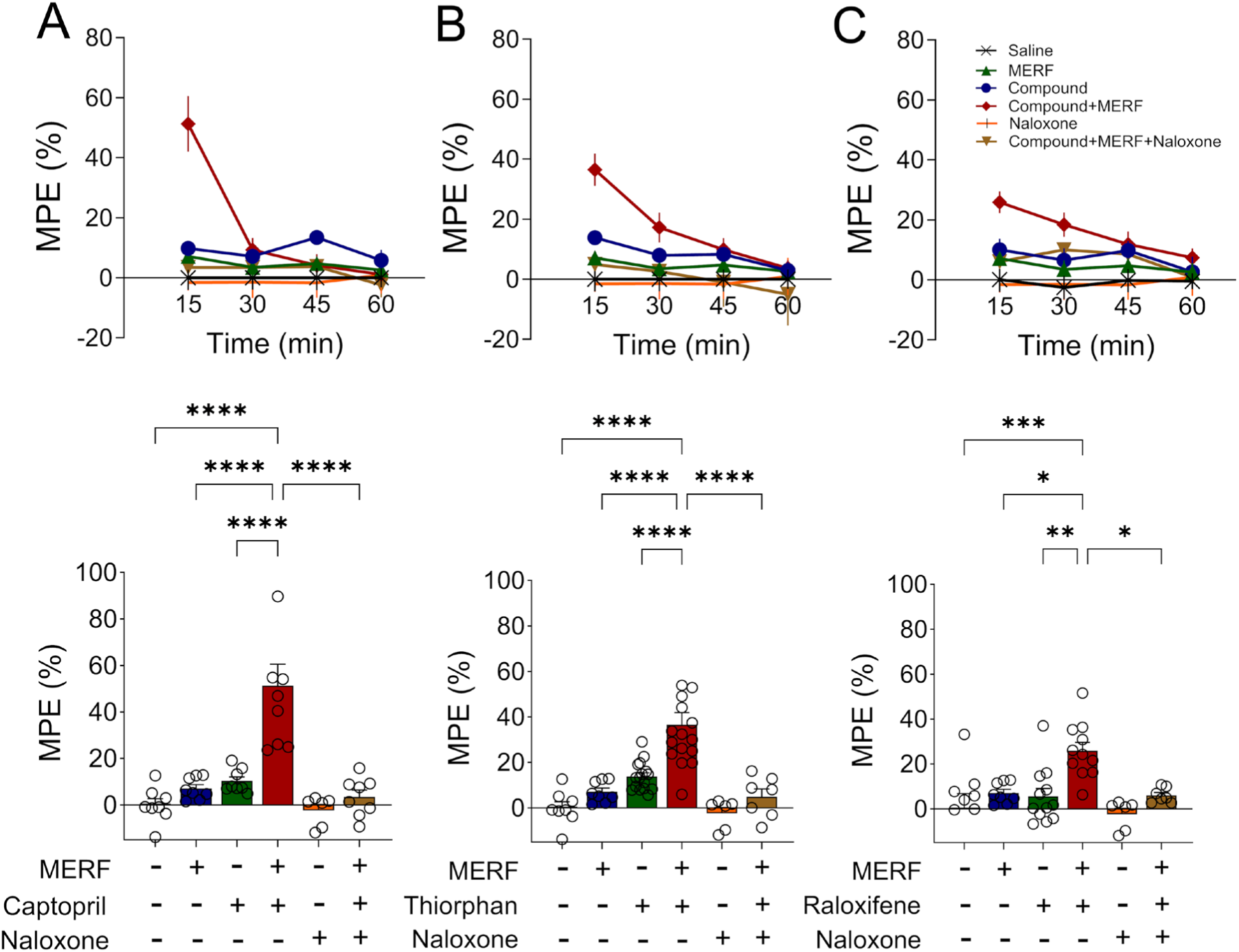
Assessment of the antinociceptive effect of the lead compounds by Hot plate assay. Compounds (100 μg/mouse) were administered to the mouse brain by intracerebroventricular (*i.c.v.*) route according to the designed treatment regimen described above and %MPE of the compounds was recorded (N = 6–15/group). A time-course of the antinociceptive effect of the lead compounds; captopril (A), thiorphan (B), and raloxifene (C), observed in the hot-plate assay is shown in the top panel. The Y-axis denotes % MPE and the X-axis shows the different time points. The bottom panel shows an analysis of %MPE for various treatment groups in the top panel observed at 15 minutes time point post compound administration. Data are shown as the mean ± SEM. One-way ANOVA followed by Sidak’s post-hoc multiple comparison test was used to determine the statistical significance (* *p* < 0.05, *** *p* < 0.001, **** *p* < 0.0001).

### Domain selectivity analysis of ACE inhibition offered by the lead candidates

Collective results of the enzymatic screening and *in vivo* experiments suggest that the *in vivo* analgesic effect of the compounds is mediated through inhibition of ACE enkephalinase activity. Somatic ACE is a membrane protein whose N- and C-termini contain zinc-dependent catalytic protease domains. Despite considerable catalytic site sequence and structure homology, the two domains differ considerably in their substrate specificity and affinity for different ACE inhibitors. Thus, we sought to determine if the lead compounds identified through this screen display differential domain affinity. To differentiate the domain-specific effect, we employed previously reported domain-selective fluorescent substrates, *i.e.*, Abz-LFK(Dnp)-OH as the C-domain specific substrate and Abz-SDK(Dnp)P-OH as N-domain specific substrate [38]. Michaelis Menten constants were determined for both substrates and enzyme kinetic curves are plotted as shown in Figure 6A and 6B. The *K*_m_ values of the C-domain substrate (Abz-LFK(Dnp)-OH) and N-domain substrate Abz-SDK(Dnp)P-OH are 10.6 ± 0.37 μM and 52 ± 5.8 μM, respectively, against mACE. These *K*_m_ values are consistent with the values reported previously, 4-12 μM for C-domain selective substrate and 35-41 μM for N-domain selective substrate [38–40]. The calculated IC_50_s and inhibitory constants *K*_i_ of the inhibitors from the enzymatic assays with these substrates (Figure 6C and 6D) are reported in Table 2. The IC_50_ of captopril was the lowest for both domains compared to other compounds, as observed with the non-domain selective substrate. Consistent with previous reports, the inhibitory constant *K*_i_ of captopril was higher for the C-domain (3.0 ± 0.033 nM) compared to the N-domain (0.57 ± 0.09 nM), suggesting a marginally higher affinity toward the N-domain [41, 42]. Thiorphan and raloxifene were more potent against the C-domain compared to the N-domain, as suggested by IC_50_s (thiorphan: N-domain: 1300 ± 80 nM vs C-domain: 770 ± 46 nM; raloxifene, N-domain: 320000 ± 43 nM vs C-domain: 140000 ± 50 nM) and lower *K*_i_ values (thiorphan, N-domain: 710 ± 45 nM vs C-domain: 170 ± 10 nM; raloxifene, N-domain: 180000 ± 24 nM vs C-domain: 30000 ± 0.05 nM) and offering selectivity factor in the range of 4.2-6 toward C-domain. D609 appears to be non-selective for both ACE domains and showed poor efficacy in these assays. These studies show differing domain preferences displayed by captopril, thiorphan, and raloxifene.

**Figure 6.**
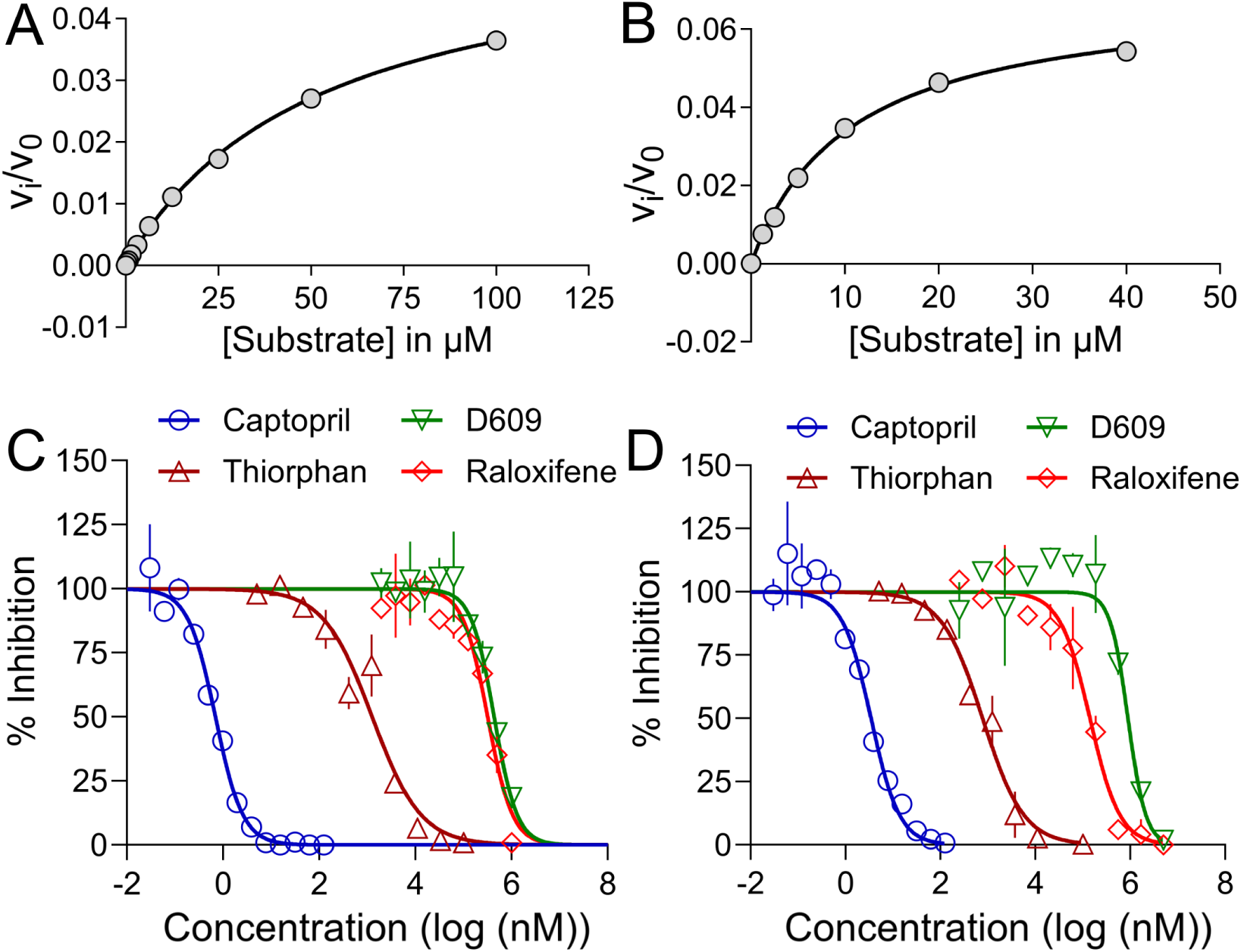
Analysis of ACE domain selectivity displayed by the lead compounds from the LOPAC screen. (A-B) The Michaelis-Menten kinetics of the ACE reaction using fluorescent peptide substrates specific for the N-domain (Abz-SDK(Dnp)P-OH, A) and C-domain (Abz-LFK(Dnp)-OH, B) of ACE. The enzymatic reaction was performed with increasing concentrations of the fluorescent substrates. The initial velocity (*V*_i_) of the reaction was calculated as described in Methods and a graph was plotted to calculate *K*_m_ values. (C-D) Dose-dependent inhibition of ACE using the domain-specific substrates (N-domain, C; C-domain, D) in the presence of increasing concentrations of the selected inhibitors — captopril (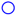), D609 (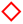), thiorphan (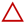), and raloxifene (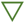).

**Table 2.** Calculated IC_50_s and *K*_i_ of the hits from N- and C-domain specific enzymatic assay.

| Compound | IC<sub>50</sub> (nM) |  | K<sub>i</sub> (nM) |  | C/N selectivity factor |
| --- | --- | --- | --- | --- | --- |
|  | N-domain | C-domain | N-domain | C-domain |  |
| Captopril | 0.57 ± 0.087 | 3.0 ± .034 | 0.57 ± 0.09 | 3.0 ± 0.033 | 0.19 |
| D609 | 430000 ± 34 | 890000 ± 60 | 240000 ± 19 | 190000 ± 13 | 1.3 |
| Thiorphan | 1300 ± 80 | 770 ± 46 | 710 ± 45 | 170 ± 10 | 4.2 |
| Raloxifene | 320000 ± 43 | 140000 ± 50 | 180000 ± 24 | 30000 ± 0.05 | 6 |

### Molecular Modeling and Docking Studies

To gain insight into the modest domain preference displayed by the lead inhibitors, we performed molecular docking studies using the available crystal structures for the N- and C-domains of ACE. X-ray crystal structure of the N-domain of ACE (PDB code 3NXQ) has been reported with a phosphinic acid-based selective inhibitor, RXP407. This PDB structure was imported for our ligand-receptor analysis as the role of the ACE N-domain was suggested previously in enkephalin degradation. Validation of the receptor grid generated for docking studies was performed by re-docking the co-crystalized ligand, and its superimposition with the co-crystalized ligand with RMSD of 0.23 Å (Supporting Information, Figure S3). Based on RXP407 binding with ACE, the active site is divided into four subsites (S2, S1, S1’, S2’), and contains Zn coordinating residues along with a putative S3 binding pocket [43]. These subsites are formed by residues that are involved in ACE catalytic process. The S2 subsites consists of Arg381, Tyr369, and the S1 subsite consists of Ala334, Asn494, Thr496 [44]. The S1’ subsite consists of His331, Tyr358, His491 and Tyr498, while His491, and Gln259 residues in the S2’ subsite. Our molecular docking analysis revealed that the lead compounds also orient themselves similarly to RXP407 and display interactions with these active site residues. The aliphatic thiol in captopril presents a Zn binding group [41]. Captopril displays hydrogen bond interactions with residues His331/491 in the S1’, and Tyr498/501 in the S2’ pockets, the residues important for enzyme catalysis (Figure 7A-B). Our results corroborate the previously reported captopril interactions with sACE [20, 45]. A salt bridge between the proline carboxylate of captopril and Lys489, which is in the proximity to Gln259 and Tyr498 involved in enzyme catalysis, contributes toward its stronger affinity for the N-domain. These interactions collectively resulted in a docking score of -13.45 kcal/mol (Supporting Information, Table S2). Thiorphan, raloxifene, and D609 also aligned similarly to captopril in the active site. Hydrogen bond interactions were observed between the carboxyl group of the thiorphan and Glu362 and Tyr501 along with metal interaction provided by the aliphatic thiol. Amide carbonyl of thiorphan displayed H-bond with the Ala334 through active site water, while pi-pi interaction was observed between the Trp335 present in the S3 binding pocket and benzyl ring of thiorphan with proximity of 3.55 Å (Figure 7C-D). This aromatic interaction is postulated to enhance the potency of ACE inhibition, resulting in an overall docking score of -7.76 kcal/mol for thiorphan. In the case of raloxifene, side chain piperidine nitrogen formed a salt bridge with negatively charged Asp43 with the proximity of 4.07 Å. Hydrogen bonds with Asn494 in the S2 subsite and Glu362 were observed, similar to thiorphan. Additional pi-pi interactions were observed between the benzothiophene ring and His365, as well as between the benzoyl ring and Phe490 (Figure 7E-F). These interactions contributed to a docking score of -6.43 kcal/mol for raloxifene (Supporting Information, Table S2). Although D609 was also positioned similarly in the active site pocket, many of the interactions were missing due to the overall smaller molecular size. Similar to captopril, thiocarbonate of D609 also formed a hydrogen bond with Glu362, resulting in a docking score of -3.09 kcal/mol, the lowest binding affinity among the inhibitors studied (Figure 7G-H). Glu362 is a part of the consensus sequence coordinating Zn and is involved in enzyme catalysis [46]. Interestingly, all the lead compounds were found to interact with this conserved residue showing the ACE inhibition potency of the ligands. Overall, the binding energy scores obtained from the docking corroborated the experimental findings for the ACE inhibition efficacy of the lead compounds. Since some of the lead compounds showed a trend toward selectivity for the C-domain in our *in vitro* enzymatic assays, an attempt was made to validate those results using this computational approach. For this purpose, the C-Domain specific crystal structure (PDB ID:2OC2) of the ACE enzyme was selected for the docking, and binding energy was calculated. (Supporting Information, Figure S4). Supplementary Table 2 shows the comparison between scores of the N-domain and C-domain selective binding free energy observed during the virtual docking. Comparable scores in both domains indicate the absence of noticeable domain-specific differences in binding interactions and energy for these inhibitors and corroborate with the experimental domain selectivity analysis.

**Figure 7.**
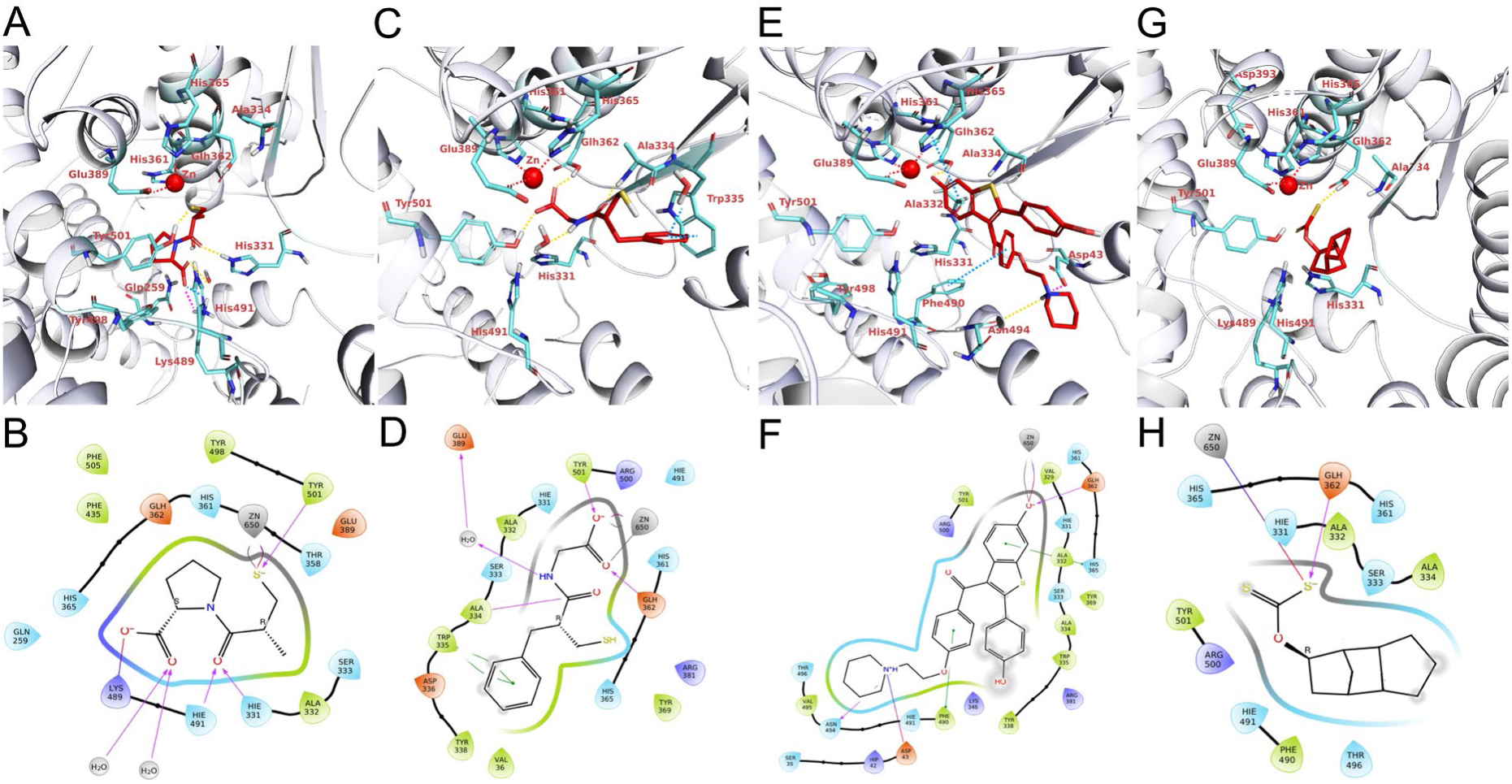
Molecular docking analysis of the binding of the lead compounds with ACE active site. The upper panel shows 3D images depicting the alignments of the lead compounds (captopril, A; thiorphan, C; raloxifene, E; and D609, G) with active site Zn (II) and the surrounding catalytic residues within the ACE active site. The lower panel includes the 2D diagrams showing the interactions of the lead compounds with the Zn (II) and the surrounding residues. The aliphatic thiol of captopril and carboxylate of thiorphan were found to be in proximity to Zn. Similar metal binding was offered by benzothiophene and thiocarbonate present in raloxifene and D609, respectively. Additionally, pi-pi and electrostatic interactions of thiorphan and raloxifene contributed favorably toward their docking scores.

## Discussion

With the ongoing opioid epidemic, approaches toward increasing endogenous neuropeptide levels by inhibition of enkephalinases have drawn significant attention. Such strategies have shown benefits in preclinical studies by inducing anti-nociceptive effect without abuse or overdose liability [47, 48]. The relationship between ACE and enkephalins, specifically in the degradation of MERF, was recently reported by us [3] and has been suggested by a few prior studies [15, 18]. Due to its carboxypeptidase A-like activity, ACE also participates in the kallikrein-kinin system, where it catalyzes the inactivation of kinins such as kallidin and bradykinin that bind to B1 and B2 receptors, known to modulate pain perception. Indeed, modest antinociceptive effects of clinically used ACE inhibitors have been observed in preclinical and clinical studies [2]. Despite these observations, efforts to identify and develop central ACE inhibitors for understanding their potential as non-opioid analgesics with better safety profiles are rather limited [49, 50]. This study provides a proof-of-concept of this rationale by utilizing high-throughput screening of a pilot library of pharmacologically active compounds (LOPAC) to identify chemically diverse leads and further confirmation of their effect on endogenous opioid signaling. The modest activities of the leads suggest the requirement of further structural explorations to improve ACE inhibition and CNS-specific effects.

A key component of the approach involved the establishment of an optimized fluorescence-based ACE enzymatic assay suitable for HTS and compatible with a 384-well plate format. A non-domain specific, FRET substrate Mca-RPPGFSAFK-(Dnp)- OH, based on bradykinin sequence, with a putative cleavage site between phenylalanine and serine, was employed for fluorescence measurement [51, 52]. The optimization of the assay involved the selection of EDTA as a reaction quencher and confirmation of the tolerance of the reaction conditions to DMSO concentration employed in the assay to enable endpoint measurement during this high-throughput screen. Overall, the assay provided an excellent Z’-factor of 0.76, confirming the quality of the assay to identify true hits with high reproducibility. Our selection of the hits was then guided by inhibitory potency, literature suggesting potential CNS activity, and modulation of pain. After the exclusion of general metal chelators, thiorphan, D609, and raloxifene were selected for further confirmatory studies. Reported as a neprilysin inhibitor, thiorphan has also been studied for the regulation of endogenous opioid-induced changes in emotional and behavioral response [53, 54]. Approved for osteoporosis, an estrogen receptor modulator raloxifene has been reported for its effect on nociception and potential CNS effect [55]. D609 inhibits phosphatidyl choline-specific phospholipase C (PC-PLC) with chelation of Zn and exhibits a myriad of pharmacological actions, including antioxidant, anti-inflammatory, and neuroprotective activities [56]. *In vitro* dose-dependence studies using fluorescence and LC-MS/MS-based methods confirmed their ACE inhibitory potency and offered IC_50_ in the nanomolar to micromolar range. Although these inhibitors were less potent than captopril, these molecules are physiologically safe, pharmacologically active, and provide opportunities for further structural explorations in future studies.

To examine the utility of ACE inhibition in enhancing endogenous enkephalin levels, we determined the effect of these leads on enkephalin release from coronal brain slices. Our previous study displayed specific enhancement of MERF by captopril and suggested the role of ACE in the degradation of this heptapeptide. Mass spectrometry-based measurement of the endogenous opioid peptides revealed the selective effect of raloxifene on MERF compared to its effect on the degradation of classical Met- and Leu-enkephalins. Thiorphan did not offer such distinction in these enkephalins, possibly due to contribution from neprilysin and potentially other non-ACE specific actions of thiorphan [18]. This data displays the ACE-specific effects of raloxifene in the regulation of MERF release.

Experiments in acute nociceptive assays further corroborated the *in vitro* results. Compound administration as single agents attenuated pain perception in the tail-flick and hot plate models, suggesting regulation of endogenous opioid neuropeptides, partly due to their inhibition of ACE enkephalinase activity. This finding was further supported by significant potentiation of analgesic effect when the compounds were administered in combination with MERF. Given the absence of analgesia produced by MERF due to its rapid degradation, the data suggests that these compounds at the concentrations given were able to protect exogenous MERF from degradation by enkephalinases, possibly involving ACE, and producing an analgesic response in this model. The increase in the %MPE was completely inhibited by naloxone, confirming the activation of the opioid receptor due to the enhancement of enkephalin levels. This result also raises the enticing prospect of utilizing centrally-active ACE inhibitors as adjuvants to exogenous opioid receptor agonists, such as morphine or fentanyl to achieve substantial reduction in the latter’s effective dose and improving or eliminating their reward potential. A few reports suggest the analgesic potential of captopril due to its effect on enkephalins resulting from ACE inhibition [15, 17]. Further, *i.c.v.* administration of thiorphan has also been shown to induce analgesic response in rodents by inhibiting enkephalinases in the brain [57, 58]. Other strategies for boosting endogenous opioids such as the development of Dual ENKephalinase Inhibitors (DENKIs) inhibiting enzymes aminopeptidase-N and neprilysin, have shown promise in preclinical testing [47, 59]. We have also reported upon the analgesic potential of aminopeptidase-N and puromycin sensitive aminopeptidase [57, 60], both with a documented role in enkephalin degradation. Such strategies are expected to strengthen the body’s defense against pain perception, preventing overstimulation of opioid receptors, and thus consequent development of addiction and dependence, presenting much-needed non-opioid analgesics for the management of pain.

Somatic ACE consists of two extracellular N- and C-domains with conserved Zn^2+^ binding motif, which are responsible for differential physiological functions. MERF degradation function of ACE is suggested to be related to its N-domain activity [24]. Domain-selectivity analysis of our hits displayed a slight preference of captopril toward the N-domain, while thiorphan and raloxifene marginally preferred the C-domain. It is reported that the smallest molecular size of captopril lacking a bulkier P1 group, which ensures the C-domain selectivity, could be responsible for its slight N-domain preference [44, 61]. The highest C-domain binding selectivity was observed in the case of raloxifene which could be due to its bulkier molecular size. Differences in physiological factors like size, shape, charge, and hydrophobicity of compound and protein complex are considered as responsible factors for the varying degree of the C/N domain selectivity [38]. *In silico* studies suggest that the lead compounds aligned and docked near the active site Zn (II) and catalytic residues of ACE, similar to captopril. Corresponding with the results of *in vitro* and *in vivo* studies, captopril showed the least binding energy, *i.e.,* strongest binding to ACE, followed by thiorphan, raloxifene, and D609, respectively. Despite this trend, we did not observe domain-specific binding energy differences among the hits. Although docking parameters may not provide sufficient insight into the domain-specific effects of these inhibitors, a comparison of active site binding interactions with RXP407, an N-domain selective ACE inhibitor, could be valuable for future structural explorations. For example, a small substituent at the P1 site, and a polar interaction offered by the P2 residue could be valuable for achieving N-domain preference. By targeting such a specific ACE domain could minimize the side effects and improve the therapeutic window of such inhibitors [21, 41].

## Conclusion

This study describes a high-throughput screening approach to identify chemically diverse ACE inhibitors, suitable for further structural explorations. The rationale for these experiments was based on our recent finding demonstrating the role of ACE in endogenous opioid signaling in striatal brain tissue. The optimized ACE enzymatic assay suitable for the HTS application identified primary hits, which were validated in our secondary enzymatic and biochemical assays, *in vivo* efficacy studies, and computational analysis. Thiorphan and raloxifene emerged as potential leads, with raloxifene offering direct ACE-specific effects. These investigations confirmed the enhancement of MERF signaling by ACE inhibition and the potential application of this approach in the development of safer analgesics, useful as single agents or adjuvants to opioids. Collectively, the *in vitro* and *in vivo* pharmacological results along with *in silico* understanding of ACE binding have provided a rationale for future structural exploration studies to obtain the next generation of domain-selective ACE inhibitors with better efficacy and safety profiles. The current study has some limitations as the lead compounds have modest ACE inhibitory activity compared to captopril. Further, the *in vivo* effects of the inhibitors were studied by direct brain administration and need to be further evaluated by systemic administration after structural optimization to determine the viability of this concept to derive clinically useful analgesics. Reinforcing the body’s innate defense against pain by enhancing endogenous opioid neuropeptides is an attractive strategy for the development of potent and safer non-opioid analgesics. Such investigations are expected to provide an effective address to the growing burden of the opioid epidemic by providing effective adjuvants to enable opioid dose reduction without compromising their efficacy. Efforts are currently underway for structural optimization of the hits to incorporate domain-selectivity and potent ACE inhibitory properties to further explore this promising avenue for the clinical management of pain.

## Methods

### Reagents for HTS assay

Recombinant mouse ACE1 enzyme (Cat# 1513-ZN) and fluorogenic peptide substrate Mca-RPPGFSAFK(Dnp)-OH (Cat# ES005) were purchased from R&D Systems. MES (Cat# M-8250), substrates Abz-SDK(Dnp)-OH (Cat# A5730), Abz-LFK(Dnp)-OH (Cat# A5855), and the high throughput compound library (LOPAC, Library of Pharmacologically Active Compounds) (Cat# LO1280) were obtained from Sigma Aldrich. MERF peptide (Cat# P000775) was purchased from AAPPTEC. (2-hydroxypropyl)-β-cyclodextrin (Cat# H107) was purchased from the Millipore Sigma. Lead compounds for validation (captopril, thiorphan, raloxifene hydrochloride, and D609 potassium) were acquired from Cayman chemicals.

### Animals

All animal experiments were approved by the Institutional Animal Care and Use Committee (IACUC), University of Minnesota and were performed in compliance with the national ethics guidelines. All the possible attempts were made to minimize the animal distress and suffering. Animals were housed in groups of four mice per cage under controlled conditions with proper access to food and water *ad libitum*.

### ACE Enzymatic assay

Compound dissolved in DMSO or DMSO control (1 µL) was pre-incubated with 15 µL mouse ACE (mACE) enzyme diluted to 0.5 ng/µL in assay buffer (50 mM MES pH 6.5, 0.0025% IGEPAL and 1 mM TCEP) in a black 384 well plate for 15 minutes at room temperature. The reaction was initiated by the addition of 14 µL fluorogenic substrate (Mca-RPPGFSAFK(Dnp)-OH) diluted to 21.4 µM in the assay buffer. After incubation for 30 minutes at room temperature, the reaction was quenched by addition of 10 µL of 1N HCl or 4 mM EDTA. The fluorescence intensity was measured using SpectraMax M5e (Molecular Devices) at an excitation wavelength of 320 nm and an emission wavelength of 405 nm. The percentage inhibition was calculated after background subtraction as (FL_positive_ – FL_negative_)/FL_positive_ x 100. Compounds with percent inhibition lower than -50% were excluded as being autofluorescent. The threshold for identifying hits was calculated as the mean percent inhibition plus three standard deviations. The Z’ score to indicate the quality of the assay was calculated as:

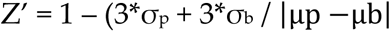

wherein, σ_p_ and σ_b_ are the standard deviation of the assay positive and negative (blank) control, respectively, and |µp −µb| is the absolute value of the difference between the means of positive and negative controls.

Further, dose–response studies of the identified lead compounds were performed using the same enzymatic assay protocol using various concentrations of compounds. IC_50_ values were calculated using the equation: (*V*_i0_-*V*_ij_)/*V*_i0_, where*V*_i0_ represents initial velocity without inhibitor and *V*_ij_ represents initial velocity observed with the selected LOPAC hit.

Initial velocities were calculated based on the linearity of the curve, typically using fluorescence data collected within the first 30 minutes. *K*_m_ and *V*_max_ values from three individual experiments were calculated using a nonlinear Michaelis-Menten fitting of initial velocities as a function of fluorogenic peptide concentration using GraphPad Prism software (version 10.0.0).

### Secondary validation of leads by LC-MS/MS analysis

The ACE inhibition assay for selected hits was performed as described in the Dose-Response Studies. Captopril (5 mM) was used as a positive control and DMSO-only reaction served as a negative control. The enzymatic reaction (30 μL) was allowed to proceed for 30 min and diluted before dilution with the assay buffer (10 μL) and further dilution with 120 μL of acetonitrile (ACN). The reaction mixture was then centrifuged at 15000 rpm at room temperature and the supernatant collected was diluted with 40% ACN to achieve a final concentration of the substrate peptide as 40 nM. The reaction mixture (1 μL) was analyzed on a TSQ Quantiva Triple Quadrupole (Thermo Scientific) in positive nanospray ionization mode using an Acclaim C8 column stabilized at room temperature (5 µm, 120 Å, 100 mm x 75 µm; Thermo Scientific, MA) and 5 mM ammonium acetate as the mobile phase to quantify the peptide [62]. Multi-step gradient elution was carried out with a flow rate of 1 μL and 0.3 μL min^-1^ and a total run time of 26 minutes. Selective reaction monitoring (SRM) was conducted by selecting transitions specific for the fluorogenic peptide [m/z 694.79 → 313.11, 694.79 → 858.37 and 694.79 → 929.41]. The optimized peptide-dependent operating parameters are as follows: static spray voltage with positive ion spray voltage 2 kV and negative ion spray voltage 0.6 kV, collision energy 25.9 V, and ion transfer tube temperature 350 °C. The residual peptide concentrations after ACE catalysis were recorded and used to calculate the IC_50_ values of the lead compounds as mentioned above.

### Mass Spectrometry-based quantification of endogenous enkephalin levels

Endogenous enkephalin release was evoked from coronal slices of the striatum, taken from C57/BL6 WT mice, by individually submerging each slice (300 µm thick) in 100 µL of aCSF containing 50 mM KCl for 20 minutes. Slices were submerged in the presence or absence of lead compounds. The final concentrations of lead compounds were 10 µM for captopril and thiorphan, and 500 µM for raloxifene and D609. Endogenous enkephalin levels in the extracellular fluid of each sample were analyzed by LC-MS/MS, as described previously [3]. Briefly, samples underwent solid phase extraction using C-18 stage tips (SP301, Thermo Scientific) and were concentrated *in vacuo*. Concentrated samples were reconstituted in 12 µL of water:acetonitrile:formic acid (98:2:0.1), and 5 µL of sample was injected onto an analytical C-18 reverse phase column (Phenomenex, Torrance, CA) using a gradient elution with 0.1% FA in water and 0.1% FA in ACN as mobile phase at 1 µL/min flow rate. Previously established mass transitions [3] for the enkephalin peptides were used for detection on a TSQ Quantiva Triple Quadrupole (Thermo Scientific) in positive nanospray ionization mode. Peaks were analyzed using Skyline and manually inspected for accuracy.

### Acute analgesia measurement: Tail-Flick Assay

Analgesic efficacy of the lead ACE inhibitors was assessed by a tail-flick assay using Columbus Instruments tail-flick analgesiometer. Stock solutions of captopril and D609 (10 mg/mL) were prepared in saline, while thiorphan was dissolved in 5% DMSO in saline. Raloxifene was dissolved in saline containing 5% DMSO and 20% (2-hydroxypropyl)-β-cyclodextrin. MERF was combined with the inhibitors at a final concentration of 5 mg/mL prior to injections. Eight-week-old male ICR-CD1 mice (weight: 22-26 gm) (Charles River Laboratories) were used in these experiments. To ensure consistency, the intensity of the heating beam was meticulously calibrated to elicit a baseline response time of 2-4 seconds. Mice were injected with the lead compounds intracerebroventricularly (*i.c.v.*) at a dose of 100 µg/mouse, either alone or in combination with MERF (50 µg/mouse) under isoflurane anesthesia, and the nociceptive response was recorded as described previously [57, 60]. Briefly, mice were positioned with the tail tip 20-30 mm from the heat source and the latency to withdraw a tail was recorded. For experiments with naloxone, subcutaneous naloxone (5 mg/kg in water) was injected 5 minutes after the *i.c.v.* injection of compounds. A cut-off of 10 s for tail withdrawal was used to prevent tissue injury.

### Acute analgesia measurement: Hot Plate Assay

Hot plate assay of the lead compounds was performed as described previously with slight modifications [57, 60], Briefly, mice were kept on the platform of the cylindrical glass bath pre-heated at 55 °C, after compound administration as described in the tail-flick assay. Latency to shake, lick, or lift the hind paw or jump from the platform was recorded at designated time points. To avoid tissue injury, a cut-off time of 60 s was selected while recording the response.

Analgesic response by tail-flick and hot plate assay are expressed in the form of percentage maximum possible effect (%MPE). The %MPE was calculated using the equation: %MPE = (Test−Baseline)/(Cut-off −Baseline) × 100, where, Test is the reaction time observed after compound administration, Baseline is the latency prior to the treatment, and Cut-off is the maximum recording time for ending the test, which is set at 10 and 60 seconds for the tail flick and hot plate assays, respectively.

### ACE domain-selectivity analysis of the lead candidates

Dose-dependent inhibition of each domain of ACE by the lead compounds was performed by following the enzymatic assay protocol discussed above with slight modifications. Briefly, mACE was incubated with previously reported domain specific substrates: Abz-SDK(Dnp)P-OH for the N-domain of mACE and Abz-LFK(Dnp)-OH for the C-domain. To test N-domain-mediated cleavage, 35 nM mACE is pre-incubated with test compounds briefly in buffer (100 mM Tris, 50 mM NaCl, 10 μM ZnCl_2_, pH 7.0) in a black 384 well plate before starting the reaction with 40 µM Abz-SDK(Dnp)P-OH. To test C-domain-mediated cleavage, 7 nM mACE is pre-incubated with test compounds briefly in buffer (50 mM HEPES, 300 mM NaCl, 10 μM ZnCl_2_, 0.0025% IGEPAL, pH 7.4) in a black 384 well plate before starting the reaction with 40 μM Abz-LFK(Dnp)-OH. For both substrates, the plate is incubated for 30 minutes at 37°C then read kinetically for 1.5 h at 37°C on a SpectraMax M5e spectrophotometer at ex/em = 320/420 nm. Fluorescence measurements were used to plot the curve of % maximal activity vs log concentration of the substrates and IC_50_ and *K*_i_ values of each compound were calculated for each domain using GraphPad Prism software (version 10.0.0 (153)). Enzyme reactions containing different concentrations of the substrates were performed to determine *K*_m_ of both domain-specific substrates.

### Virtual screening and molecular docking of the lead compounds

Molecular modeling was performed using the Schrödinger Maestro 2022-2 by following the protocol described by Singh et al. [60] with slight modifications. The crystal structure of ACE (PDB: 3NXQ) complex with a synthetic ligand, RXP407 was used to study the interactions of the lead compounds. A protein preparation wizard was utilized to prepare the protein structure for the docking. Protein structure was preprocessed by adding the missing hydrogen atoms and removing the water molecules beyond the radii of 5 Å. Missing side chain and loop residues were incorporated using the Prime tool, while PROPKA at pH 7.0 was used to optimize the hydrogen bonds assignment. Protein structure was further minimized using the OPLS4 force field. To confine the docking space on the receptor for the ligands, the grid was prepared using the receptor grid generation tool. The endogenous ligand was considered as a center point and a receptor grid was generated, which encompasses all active site amino acid residues within a 15 Å^3^ box. Coordinates for the receptor grid generated are X: -1.53 Å, Y: -16.64 Å, and Z: -22.14 Å in the receptor. By applying partial charge cut-off 0.25 and van der Waals radii of nonpolar atoms in the receptor, the grid’s flexibility was ensured. After the protein preparation, ligands were prepared in order to make them suitable for docking using LigPrep tool of Maestro. Prior to ligand preparation, lead compounds captopril, thiorphan, D609, and raloxifene were sketched using the 2D-sketcher. Ionization was carried out using Epik ionizer and ligands conformers with different metal ion states were obtained at a pH range of 7 ± 2. Ligands prepared were subjected to docking with the prepared receptor protein. Docking was performed using the Extra Precision (XP) mode of the Glide module. During docking, all the prepared ligands were considered flexible entities with the assignment of intramolecular hydrogen bonding. Poses generated after the docking were subjected to post-docking minimization and strain correction, and the binding energy score in the form of the dock score was noted for each ligand.

## Author Contributions

Conceptualization, S.S.M. and P.E.R.; experiments, P.D., J.W., and F.H.; data curation and analysis, P.D., J.W., F.H., P.E.R., and S.S.M.; funding acquisition, S.S.M. and P.E.R.; writing—original draft preparation, P.D.; writing—review and editing, S.S.M., P.E.R., F.H., and J.W.; supervision, S.S.M. and P.E.R. All authors have read and agreed to the published version of the manuscript.

## Institutional Review Board Statement

The study was conducted in accordance with the Declaration of Helsinki and approved by the Institutional Animal Care and Use Committee (IACUC) at the University of Minnesota (protocol ID 2204-39921A; date of last approval August 20, 2024). All applicable ethical standards required by the University of Minnesota, IACUC were followed.

## Data Availability Statement

The data presented in this study, such as raw data or original images, are contained in the article and Supplementary Materials. The chemical compound is available upon request from the corresponding author.

## Acknowledgments

The authors thank Yingchun Zhao and Linda von Weymarn from the Analytical Biochemistry Shared Resource of the Masonic Cancer Center at the University of Minnesota for technical help with the mass spectrometry experiments. We also thank Dr. Rohit Singh for helpful discussions related to the molecular docking studies. The graphical abstract and figure schematic were created with BioRender.com.

