## Supporting Information for "Exploiting the Angiotensin-Converting Enzyme Pathway to Augment Endogenous Opioid Signaling"

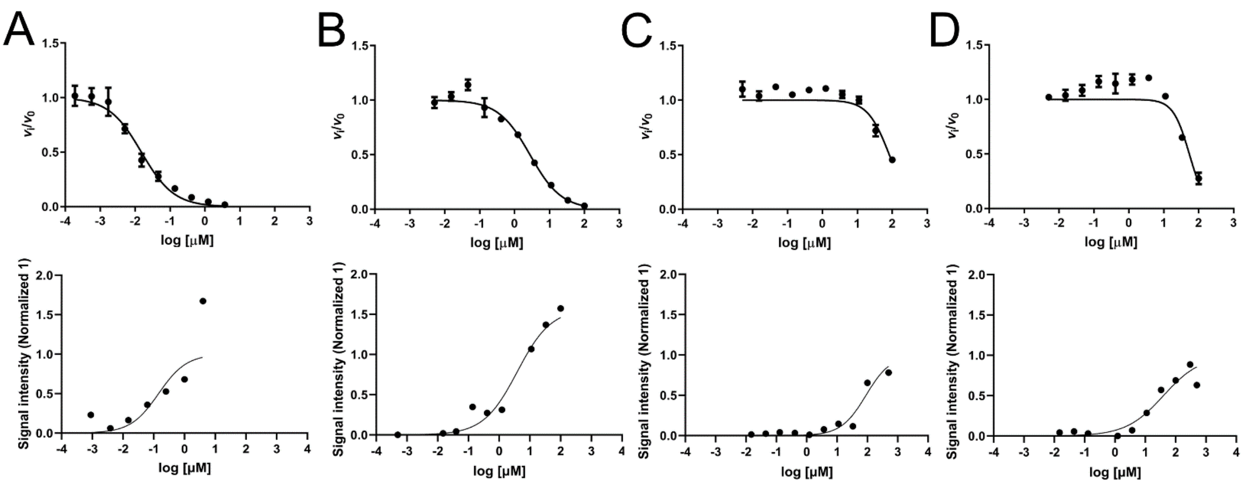


**Fig. S1.** Individual dose response curves of the lead compounds obtained from the ACE enzymatic assay using FRET assay (upper panel) and from the LC-MS/MS-based validation (lower panel); (A) captopril, (B) thiorphan, (C) raloxifene, (D) D609. Non-linear sigmoidal curve fitting was utilized to calculate $IC_{50}$ values.

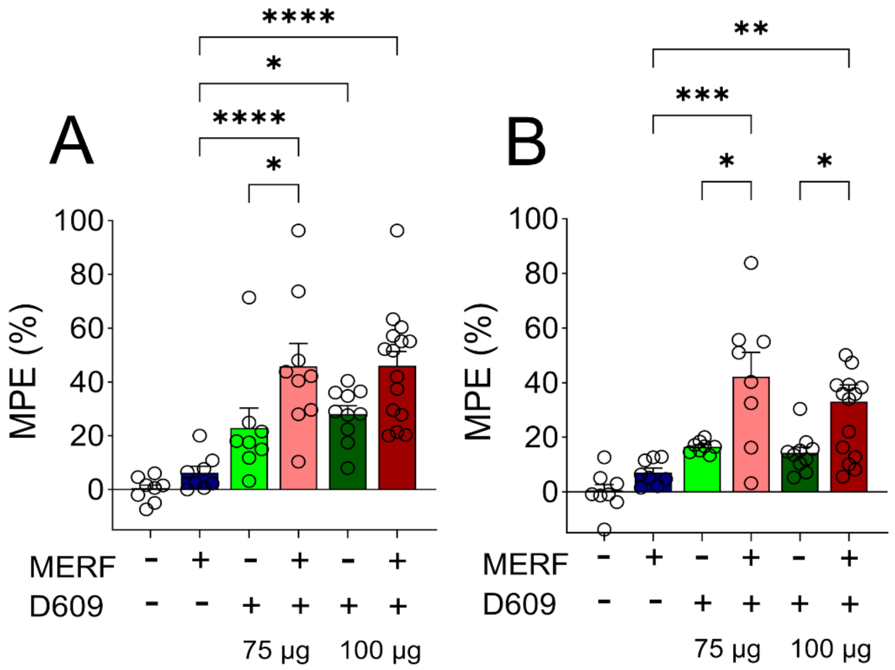


**Fig. S2.** Assessment of the antinociceptive effect of D609 at the doses of 100 µg/animal and 75 µg/animal by tail-flick (A) and hot plate (B) assays. D609 was administered directly to the mouse brain by intracerebroventricular (*i.c.v.*) route at the indicated dose, either alone or in combination with the MERF (50 µg/mouse) and the maximum possible effect of the compounds was recorded (N = 6–15/group). The bar graphs show the comparison between the anti-nociceptive effect (%MPE ± SEM) for various treatment groups at 15 minutes after compound administration. Data are shown as the mean ± SEM. Statistical significance was examined by a one-way ANOVA with Sidak's post-hoc multiple comparison test (* $p < 0.05$, *** $p < 0.001$, **** $p < 0.0001$).

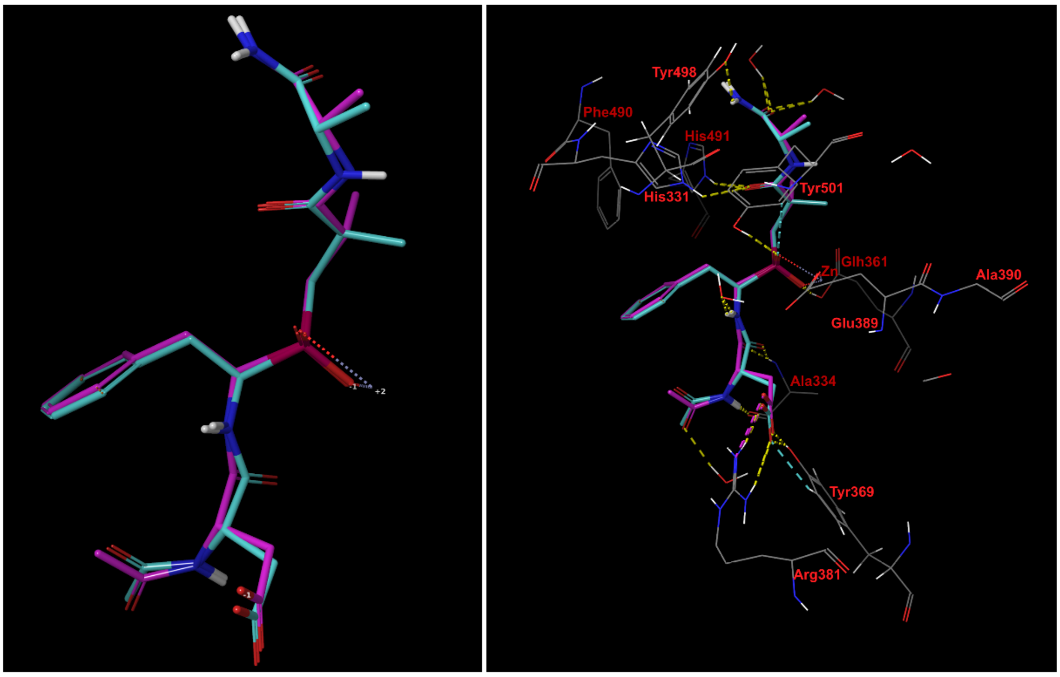


**Fig. S3.** Validation of the docking parameters by redocking the co-crystalized ligand (RXP407) using N-domain specific ACE crystal structure (PDB ID: 3NXQ). Left image: Superimposed structures of docked ligand (pink sticks) with the co-crystalized ligand (cyan sticks), Right image: Interactions observed between docked and co-crystalized ligand with the surrounding active site residues.

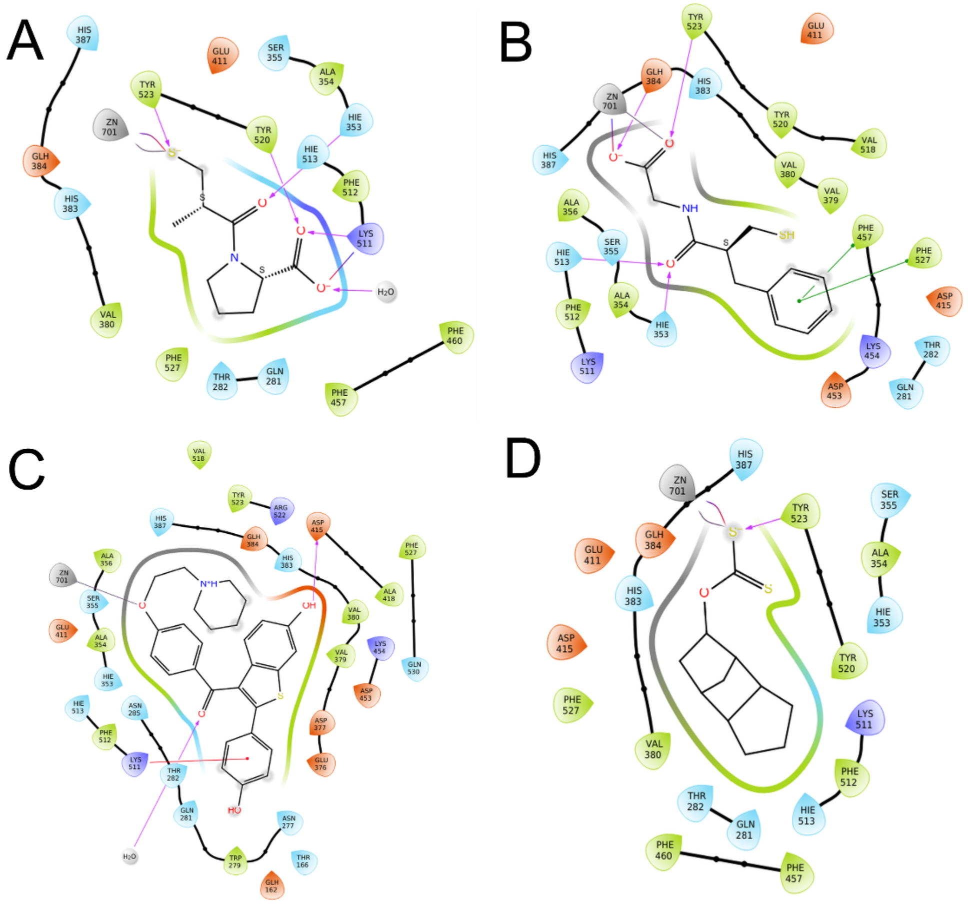


**Fig. S4.** 2D interaction diagrams of the docked lead compounds with the C-domain of ACE. (A) Captopril (B) Thiorphan (C) raloxifene and (D) D609 docked using the C-domain specific crystal structure of ACE (PDB ID: 2OC2) following the experimental protocol described in Methods.

**Table S1:** List of hits identified from the LOPAC screening based on ACE inhibitory potency using a 46% cut-off threshold, as described in Methods. Reported pharmacological actions of these compounds are referred from https://www.sigmaaldrich.com/US/en assessed on 01/02/25.

| Compound name | % Inhibition ± SD | Class/mechanism of compound |
|---|---|---|
| Enalaprilat dihydrate | 100 ± 0.01 | ACE inhibitor |
| Diethylenetriamine pentaacetic acid | 99.7 ± 0.17 | Chelator |
| SKF 89976A hydrochloride | 99.4 ± 0.93 | γ-Aminobutyric acid (GABA) transporter (GAT-1) inhibitor |
| Captopril | 98.2 ± 2.3 | ACE inhibitor |
| Trandolapril | 97.6 ± 0.24 | ACE inhibitor |
| 1, 10- Phenanthroline monohydrate | 96 ± 0.53 | Chelator |
| DL-thiorphan | 95 ± 2.9 | Neprilysin inhibitor |
| Reactive blue 2 | 89 ± 5.6 | ATP receptor antagonist |
| Fiduxosin hydrochloride | 69.4 ± 31 | α1a-adrenoceptor antagonist |
| Raloxifene hydrochloride | 64 ± 6.6 | Selective estrogen receptor modulators (SERMs) |
| D609 potassium | 61.6 ± 12 | Phosphatidylcholine (PC)-specific phospholipase C inhibitor |
| Lonidamine | 57.9 ± 35 | Glycolysis inhibitor |
| Bethanechol chloride | 51.6 ± 69 | Muscarinic agonist |
| Tetraethylthiuram disulfide | 51 ± 1.8 | Metabolic enzymes inhibitor |
| RX 821002 hydrochloride | 50 ± 21 | α2-adrenoceptor antagonist |
| IPA-3 | 48 ± 13 | Pak1 inhibitor |
| Hydrocortisone | 46.7 ± 75 | Steroid |
| Protoporphryn IX | 46 ± 2.8 | Heme and Chlorophyll precursor |
| Alinidine | 45.9 ± 19 | Negative chronotrope |

**Table S2:** Comparison of the binding energies of the lead compounds docked with the N domain (PDB ID: 3NXQ) and C domain (PDB ID: 2OC2) crystal structures of ACE.

| Name of Compound | Binding Energy Score (Dock score) (kcal/mol) | | | |
|---|---|---|---|---|
| | N domain (Zn as constraint) | N-domain (No constraint) | C domain (Zn as constraint) | C domain (No constraint) |
| Captopril | -7.32 | -13.45 | -7.22 | -11.79 |
| Thiorphan | -7.82 | -7.76 | -8.14 | -8.13 |
| Raloxifene | -5.04 | -6.43 | -1.86 | -3.43 |
| D609 | -2.62 | -3.09 | -2.33 | -2.92 |
